# Infection with *Mycobacterium tuberculosis* orchestrates the PRMT5-dependent methylation of NCOA4 to govern host ferroptosis

**DOI:** 10.1101/2025.10.23.684158

**Authors:** Smriti Sundar, Raju S Rajmani, Kithiganahalli Narayanaswamy Balaji

## Abstract

*Mycobacterium tuberculosis* (Mtb) increases availability of free iron resulting in ferroptosis of host macrophages to facilitate its survival and dissemination. A critical factor for elevated levels of labile iron is the overt accumulation of the protein nuclear receptor coactivator 4 (NCOA4), that promotes autophagic degradation of ferritin in a process termed as ferritinophagy. Here, we identify a novel post-translational modification on NCOA4 that is essential for its interaction with ferritin in iron-replete condition of Mtb-infected cells. Specifically, protein arginine methyltransferase 5 (PRMT5), confers a symmetric-dimethylation on NCOA4 that promotes ferritinophagy-mediated ferroptosis. Using loss-of-function studies, we show that PRMT5 is required for lipid peroxidation, bacterial survival, and dissemination in Mtb-infected mice. Also, overexpression of a methylation-deficient mutant of NCOA4 phenocopy depletion of PRMT5 and reduced ferritinophagy in Mtb-infected cells. Furthermore, PRMT5-mediated methylation reduces the nuclear availability of NCOA4 and impairs its co-activatory role to nuclear receptors such as vitamin D3. Thus, our findings uncover the key interaction between NCOA4 and ferritin that regulates ferroptosis and mycobacterial survival during infection. Perturbation of this interaction results in reduced Mtb loads and alleviated disease pathology.

## Introduction

Tuberculosis (TB) is characterized by dynamic metabolic interplay between *Mycobacterium tuberculosis* (Mtb) and the host wherein Mtb employs efficient nutrient retrieval strategies to survive despite the host’s efforts to deplete essential resources through nutritional immunity (*1–3*). The limited bioavailability of metal ions like iron, act as significant barriers for mycobacterial survival, and increased access to iron within its intracellular niche has been largely reported to favor Mtb growth (*4–6*). For instance, macrophages from patients of hereditary hemochromatosis, characterized by a low intracellular iron pool, impair TB growth while iron overload is associated with death from TB(*7*, *8*). Apart from fulfilling the nutritional requirements of Mtb, high levels of labile iron pool (LIP) can boost production of reactive oxygen species (ROS) and oxidize polyunsaturated fatty acids, such as those found in membrane phospholipids of the host(*9*, *10*). The uncontrolled accumulation of these lipid peroxides can irreversibly permeabilize cellular membranes and lead to ferroptosis (*11–13*).

Ferroptosis, an oxidative form of cell death triggered by cytoplasmic iron-overload and impaired antioxidant defences within the cell, has extensive implications in inflammation and immunity(*14–19*). Recent studies have identified a direct association of macrophage ferroptosis with extracellular release of Mtb, necrotic lesions, substantial tissue damage, and mycobacterial dissemination(*20–22*). This is driven by the global modulation of antioxidant responses and downregulation of GPX4 (Glutathione Peroxidase 4) and GSH (Glutathione) levels(*23–25*). However, increased availability of iron is a pre-requisite to the induction of ferroptosis and the source of such large amounts of free iron in Mtb-infected cells remains unknown.

Macrophages sequester free iron within ferritin, a multi-subunit intracellular storage protein made up of heavy and light chains to limit availability to invading pathogens(*26*, *27*). Under conditions of intracellular iron deficiency, NCOA4 (Nuclear receptor coactivator 4), the cargo receptor of ferritinophagy, selectively interacts with ferritin and MAP1LC3/LC3, initiates the phagophore formation, and recycles the bound iron to replenish the cellular LIP. During iron-replete conditions, NCOA4 undergoes HERC2 (ECT and RLD61 domain containing E3 ubiquitin protein ligase 2) mediated proteasomal degradation, thereby controlling the rate of ferritinophagy(*28*, *29*). Thus, while Mtb infection alters expression of heme catabolizing enzymes like HO-1 (Heme oxygenase-1), the free iron released into the cell can be rapidly sequestered by ferritin. Interestingly, recent findings indicate that Mtb reduces HERC2 levels, which in turn increases levels of NCOA4, and drives ferritinophagy in macrophages(*30*). Given the key role of increased intracellular iron in ferroptosis, the role of ferritinophagy, that releases large amounts of ferrous iron into the cytoplasm, must be evaluated. Additionally, overarching regulatory mechanisms such as post-translational modifications and the interactome of NCOA4 control its interaction with ferritin and may aid in ensuring that ferritinophagy is only ensued upon intracellular iron deficiency(*31*, *32*). It remains unclear how Mtb overcomes these additional regulatory machineries to induce ferritinophagy despite the iron-replete conditions within macrophages.

Protein arginine methylation, a post-translational modification mediated by PRMTs (protein arginine methyltransferases), has garnered attention in modifying key enzymes in iron metabolism and lipid peroxidation (*33–35*). PRMT5, a major type II enzyme that catalyzes the symmetric-dimethylation of arginine residues on histone/ non-histone proteins was recently identified to regulate antioxidant responses and iron homeostasis in distinct models of study (*36–38*). Importantly, PRMT5 has been previously reported by our group to assist in Mtb pathogenesis and survival by contributing to lipid accumulation. To our interest, infected mice treated with PRMT5 inhibitor also exhibited a reduction in granulomatous lesions and mycobacterial dissemination, indicative of yet-unexplored mechanisms regulated by PRMT5(*39*). This encouraged us to investigate the role of PRMT5 in the context of Mtb-driven ferroptosis and mycobacterial dissemination.

Our results identify a novel methylation on NCOA4 conferred by PRMT5, essential for its cytoplasmic retention and ferritinophagy. The consequent increase in free iron induces ferroptosis and aids in Mtb pathogenesis. Additionally, the methylation on NCOA4 compromises its nuclear availability and may thus undermine its function as a coactivator to various nuclear receptors, including estrogen receptor, androgen receptor, and vitamin D3 receptor. When assessed specifically with the vitamin D3 receptor, we demonstrated that the inhibition of PRMT5 could significantly improve the clearance of Mtb infection and enhance vitamin D3-induced immunity. Together, the findings highlight the potential role for NCOA4 and PRMT5 in Mtb pathogenesis and provide a framework to understand their implications as a novel target for TB therapeutics.

## Results

### PRMT5 activity regulates ferroptosis during Mtb infection

To investigate the role of PRMT5 in Mtb-induced ferroptosis, we employed an *ex vivo* model of infection in which macrophages isolated from BALB/c mice were pre-treated with EPZ015666 (PRMT5 inhibitor), infected with Mtb, and assessed for the key hallmarks of ferroptotic death– increased labile iron pool, increased oxidative stress, and accumulation of lipid peroxides. Inhibition of PRMT5 activity could significantly reduce cellular LIP (Figure 1A) and levels of ROS (Figure S1A, B) observed during Mtb infection. Subsequently, as a relative measure of lipid peroxidation, we assessed for levels of a lipid peroxidation-derived aldehyde byproduct, 4-hydroxynonenal (4-HNE). As expected, treatment with the PRMT5 inhibitor substantially reduced levels of 4HNE in Mtb-infected cells. (Figure 1B). The accumulation of lethal levels of lipid peroxides leads to an increase in cell death and allows the release of Mtb into the extracellular milieu(*20*). In accordance, PRMT5 inhibition compromised cell death and Mtb release into culture supernatants (Figure 1C). Further, as Mtb-induced ferroptosis in host cells is characterized by GPX4 depletion, we assessed for GPX4 levels post PRMT5 activity inhibition. Interestingly, treatment with PRMT5 inhibitor could not rescue GPX4 levels during Mtb infection (Figure S1C), suggesting that the inhibition of PRMT5 antagonizes ferroptosis through an independent manner. Next, we infected BALB/c mice with ∼10^3^ CFU of Mtb via the aerosol route and administered EPZ015666 (Figure 1D). When assessed at 56 days post-infection, the inhibition of PRMT5 protected the mice from TB pathology (Figure 1E, Figure S1D, E), consistent with our previous study(*39*), and resulted in a significant reduction in lung weight and calculated relative lung mass (Figure S1F). In addition to reduced mycobacterial burden in the lungs, mice treated with the PRMT5 inhibitor also had a significantly reduced Mtb burden in the spleen, and liver, indicating the role of PRMT5 in regulating mycobacterial dissemination (Figure 1F-H). Furthermore, consistent with our results of PRMT5 regulating ferroptosis during Mtb infection, these mice displayed lower lipid peroxides (MDA) levels in the lungs (Figure 1I).

**Figure 1.**
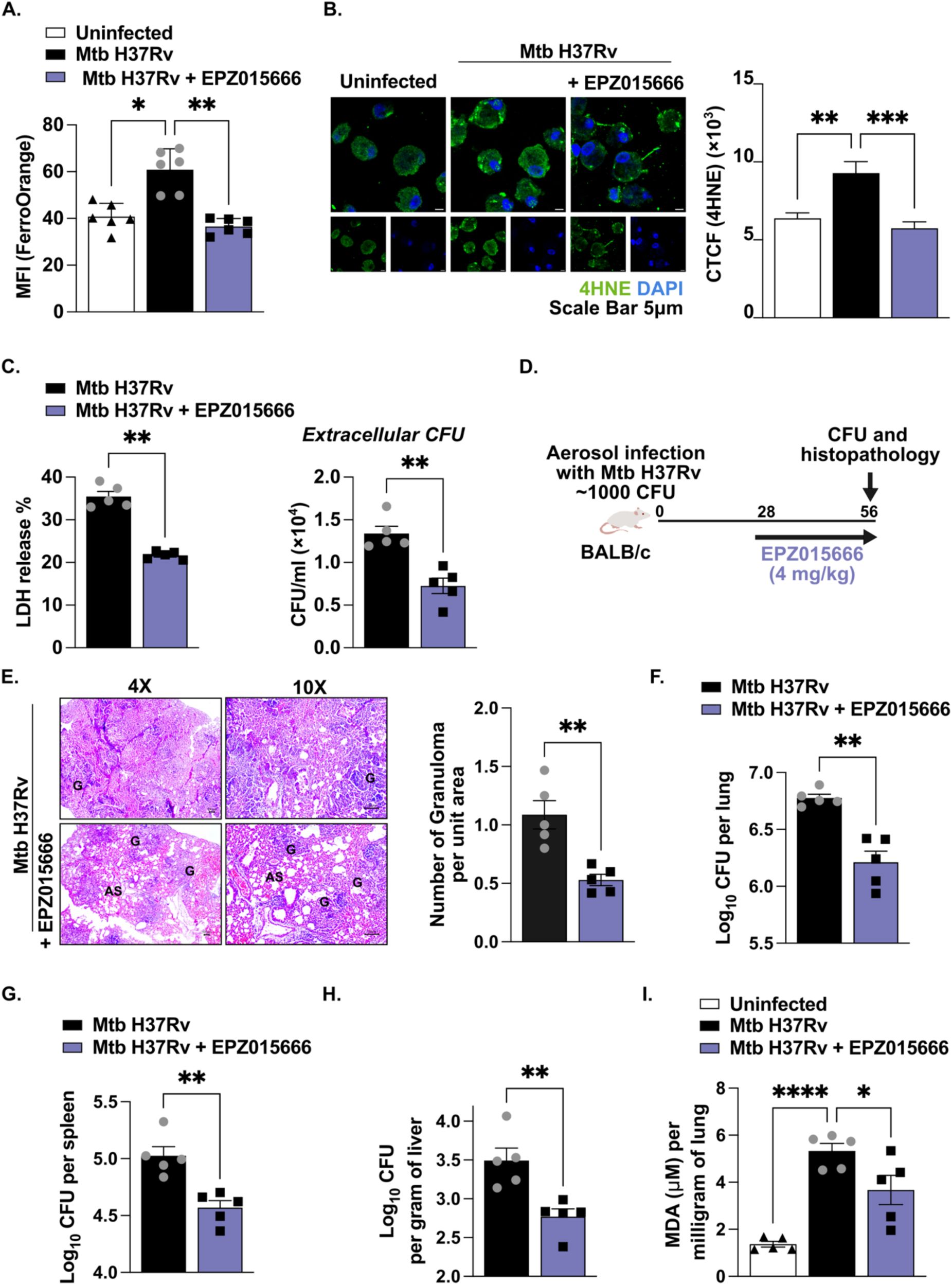
PRMT5 regulates ferroptosis during Mtb infection. (**A**) Estimation of intracellular labile ferrous iron in murine macrophages pretreated with PRMT5 inhibitor (for 1 hour) at 24 hours post infection. (**B**) Confocal images of murine macrophages pretreated with PRMT5 inhibitor (for 1 hour) stained for 4HNE (green) and nuclear staining (blue) at 24 hours post infection. Scale bars: 5 µm (left), and its quantification (right). (**C**) Assessment of necrotic cell death measured by LDH released in the supernatants from the macrophage pretreated with PRMT5 inhibitor (for 1 hour) at 4 days post infection (left). Examination of the release of live mycobacteria from necrotic cells by CFU quantification in culture supernatants of macrophages pretreated with PRMT5 inhibitor (for 1 hour) at 4 days post infection (right). (**D-J**) BALB/c mice were infected with Mtb H37Rv and treated with PRMT5 inhibitor EPZ015666 (4mg/kg) q.o.d. as indicated in (**D**) (Number of mice per group = 5) (**E**) Analysis of TB pathology (granulomatous lesions) in the lung sections with H and E staining (left) and its quantification by a pathologist in a blinded manner (right). (**F-H**) Enumeration of mycobacterial burden in the lungs (**F**), spleen (**G**), liver (**H**) of infected and PRMT5 inhibitor treated mice by plating homogenates on 7H11 plates. (**I**) Measurement of lipid peroxidation (malondialdehyde) in lung homogenates from uninfected, Mtb infected and PRMT5 inhibitor treated mice. All immunofluorescence data are representative of three independent experiments. Specific regions of the H and E-stained sections were analysed by the pathologist for evaluation of the granuloma fraction. Accordingly, the portions have been demarcated in the images; Magnification, 4X (left panel); 10X (right panel); Scale bar for H and E images, 100μm; G, granuloma; AS, alveolar space; MFI, mean fluorescence intensity; DAPI, 4’,6-diamidino-2-phenylindole; CTCF, corrected total cell fluorescence; LDH, lactose dehydrogenase; CFU, colony forming units; q.o.d., *quaque altera die*, every other day. EPZ015666 (20 μM), PRMT5 inhibitor. *, p<0.05; **, p<0.01; ****, p < 0.0001 (One-way ANOVA in A, B, I; Student’s t-test in C, E, F, G, H; GraphPad Prism 10.0).

### PRMT5 dependent induction of ferroptosis is dependent on ferritinophagy

Next, as the inhibition of PRMT5 could significantly rescue cells from Mtb-induced iron-overload, we first sought to investigate the status of the major intracellular iron storage complex, ferritin, in macrophages upon Mtb infection. We observed a robust degradation of ferritin upon Mtb infection that could be rescued upon the inhibition of autophagy (Figure S2A, B). Similarly, we observed an accumulation of the cargo receptor, NCOA4 (Figure S2C), and its depletion could also rescue levels of ferritin during Mtb infection (Figure S2D). Under homeostatic conditions, intracellular free iron is sequestered within ferritin nanocages to prevent iron-overload-associated toxicity. Ferritinophagy is subsequently activated only to mobilize iron upon requirement(*40*). In line, the addition of exogenous iron to macrophages increases the total amount of ferritin, whereas iron chelation using deferoxamine (DFO) triggers degradation, leading to a lower level of ferritin in the cells(*40*). However, this homeostatic control is lost upon Mtb infection, wherein ferritin levels are compromised despite iron availability (Figure 2A). While the increase in rate of ferritinophagy observed during Mtb infection has been attributed to accumulated levels of NCOA4 during Mtb infection, we observed that the overexpression of NCOA4 is not entirely sufficient to explain the extent of ferritin degradation observed upon Mtb infection (Figure 2B). These results together led us to hypothesize that there are additional regulators of ferritinophagy operating during Mtb infection. Immunoblotting experiments revealed that the inhibition of PRMT5 activity restored levels of ferritin heavy chain in Mtb infected cells (Figure 2C, left). Consistently, the depletion of *Prmt5* transcripts could also rescue the levels of ferritin during Mtb infection (Figure 2C, right). In addition to rescuing ferritin levels, PRMT5 inhibition could also significantly reduce ferritin’s targeting into LC3 positive autophagosomes (Figure S2E). Inhibition of PRMT5 also decreased the colocalization of ferritin with lysosomes stained with LAMP1 (Figure 2D). Consistent with these results, we also found that the reduced levels of ferritin in the lungs of mice infected with Mtb could be rescued upon treatment with the PRMT5 inhibitor (Figure 2E, S2F). Further, the depletion of NCOA4 also reduced levels of host labile iron pool (Figure S3A), and 4HNE levels during Mtb infection (Figure S3B). The depletion of NCOA4 also compromised the release of Mtb into the extracellular milieu, supporting its role in Mtb-induced ferroptosis (Figure S3C).

**Figure 2:**
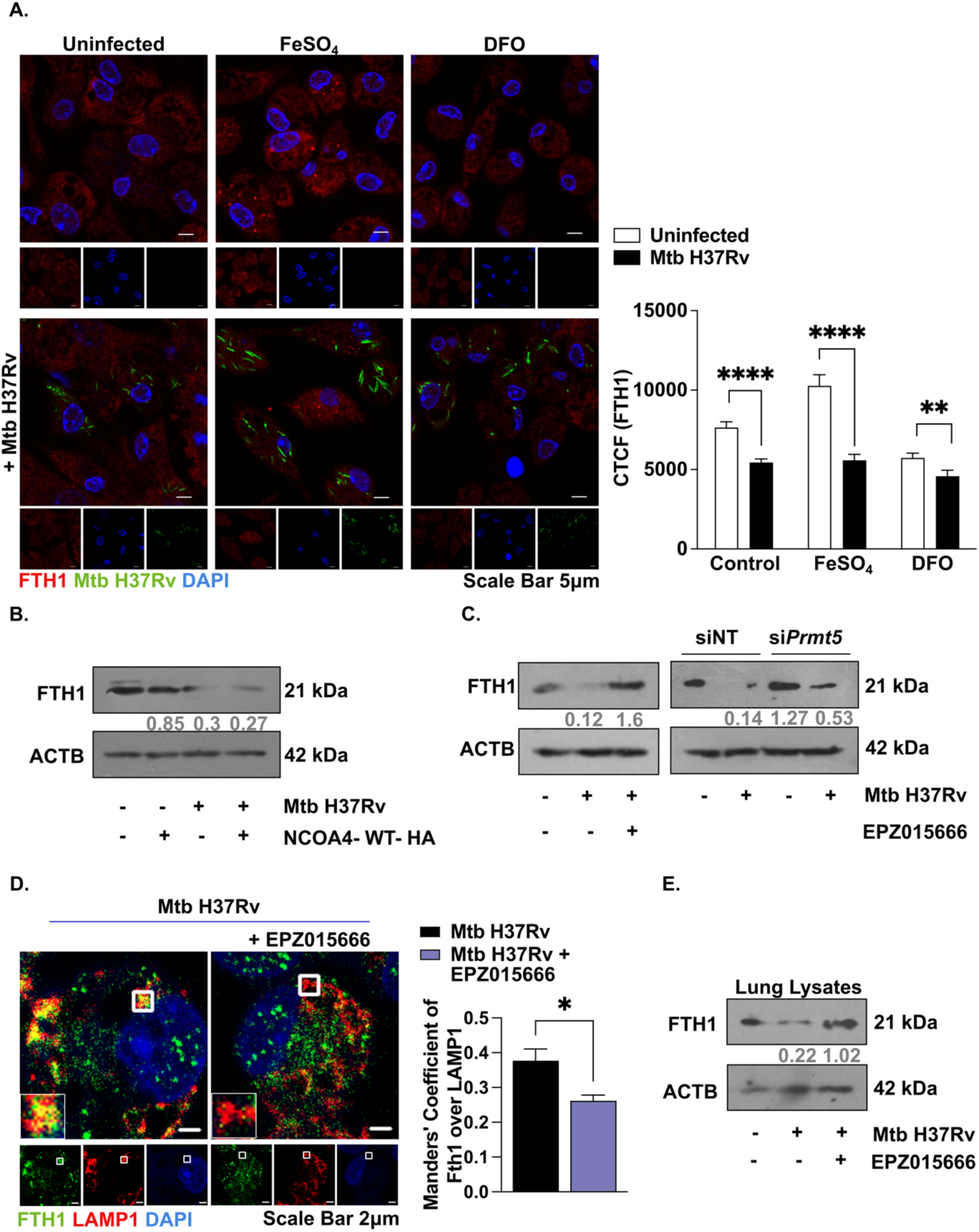
PRMT5 induces NCOA4-dependent ferritinophagy during Mtb infection. (**A**) Confocal images of murine macrophages treated with FeSO_4_ for 24 hours and then treated with DFO (iron chelator) for 1 hour, stained for ferritin heavy chain (red) and nuclear staining (blue) at 24 hours post infection. Scale bars: 5 µm (left), and its quantification (right). (**B**) Assessment of ferritin heavy chain levels in RAW 264.7 cells transfected with NCOA4 OE vectors at 24 hours post infection by immunoblotting. (**C**) Assessment of ferritin heavy chain levels in murine macrophages pretreated with PRMT5 inhibitor (for 1 hour) at 24 hours post infection (left); and in murine macrophages transfected with NT or *Prmt5* siRNAs at 24 hours post infection (right) by immunoblotting. (**D**) Confocal images of murine macrophages pretreated with PRMT5 inhibitor (for 1 hour) stained for ferritin heavy chain (green), lysosomes LAMP1 (red) and nuclear staining (blue) at 24 hours post infection. Scale bars: 2 µm (left), and its quantification (right). (**E**) BALB/c mice were infected with Mtb H37Rv and treated with PRMT5 inhibitor EPZ015666 (4mg/kg) q.o.d. Assessment of ferritin heavy chain levels in the lung homogenates of uninfected and infected mice by immunoblotting. All immunoblotting and immunofluorescence data are representative of three independent experiments. ACTB was used as a loading control. NT, non-targeting; MFI, mean fluorescence intensity; DAPI, 4’,6-diamidino-2-phenylindole; OE, over expression; CTCF, corrected total cell fluorescence; CFU, colony forming units; q.o.d., *quaque altera die*, every other day. *, p<0.05; **, p<0.01; ****, p < 0.0001 (Student’s t-test in A, D; GraphPad Prism 10.0). EPZ015666 (20 μM), PRMT5 inhibitor, FeSO_4_ (100 μM); DFO (100 μM). Below each panel, the quantification of the blots normalized to the loading control has been indicated.

### PRMT5-dependent methylation of NCOA4 is required for its interaction with ferritin

Further, to understand the mechanisms by which PRMT5 regulates NCOA4-dependent ferritinophagy, we performed a co-immunoprecipitation of NCOA4 and assessed for its interaction with ferritin. Mtb infection increased the association of NCOA4 and ferritin while PRMT5 inhibition could drastically reduce their interaction during Mtb infection (Figure 3A, S4A). Subsequently, we performed an immunostaining experiment wherein infected macrophages cultured in the presence or absence of PRMT5 inhibitor were assessed for the association between NCOA4, ferritin and LAMP1. Confocal microscopy analysis showed strong punctate structures and colocalization between NCOA4, ferritin and LAMP1 during Mtb infection which was compromised upon PRMT5 inhibition (Figure 3B, S4B). Post-translational modifications such as the symmetric di-methylation of arginine residues conferred onto proteins by PRMT5 has been identified to regulate protein-protein interaction, cellular localization and activity (*37*, *41–43*). To investigate whether NCOA4 is a bonafide substrate for PRMT5 methylation, we performed a co-immunoprecipitation of NCOA4 from cellular lysates of Mtb infected macrophages. We identified a notable increase in the symmetric di-methylation of its arginine residues (Figure S4C, D). Consistently, we found that NCOA4 interacts with PRMT5 during Mtb infection (Figure S4C) and reciprocally, co-immunoprecipitation of PRMT5 also revealed its interaction with NCOA4 (Figure S4E). Furthermore, the inhibition of PRMT5 markedly reduced the methylation of arginine residues on NCOA4 (Figure 3C, S4F). As PRMT5 preferentially methylates arginine residues present adjacent to glycine, we scanned the mouse protein NCOA4, performed a multiple sequence alignment and identified R182 as an evolutionarily conserved residue indicating it might be a critical residue methylated by PRMT5 (Figure 3D). Bioinformatic analysis could also predict a potential site for methylation at R182 of mouse NCOA4 protein (Figure S4G). To evaluate its significance, we constructed methylation-deficient mutant NCOA4-R182K and overexpressed it in macrophages. Upon infection with Mtb, the cells expressing this mutant exhibited lowered NCOA4-ferritin interaction (Figure 3E) suggesting that the methylation of NCOA4 at R182 might be necessary for its interaction with ferritin.

**Figure 3:**
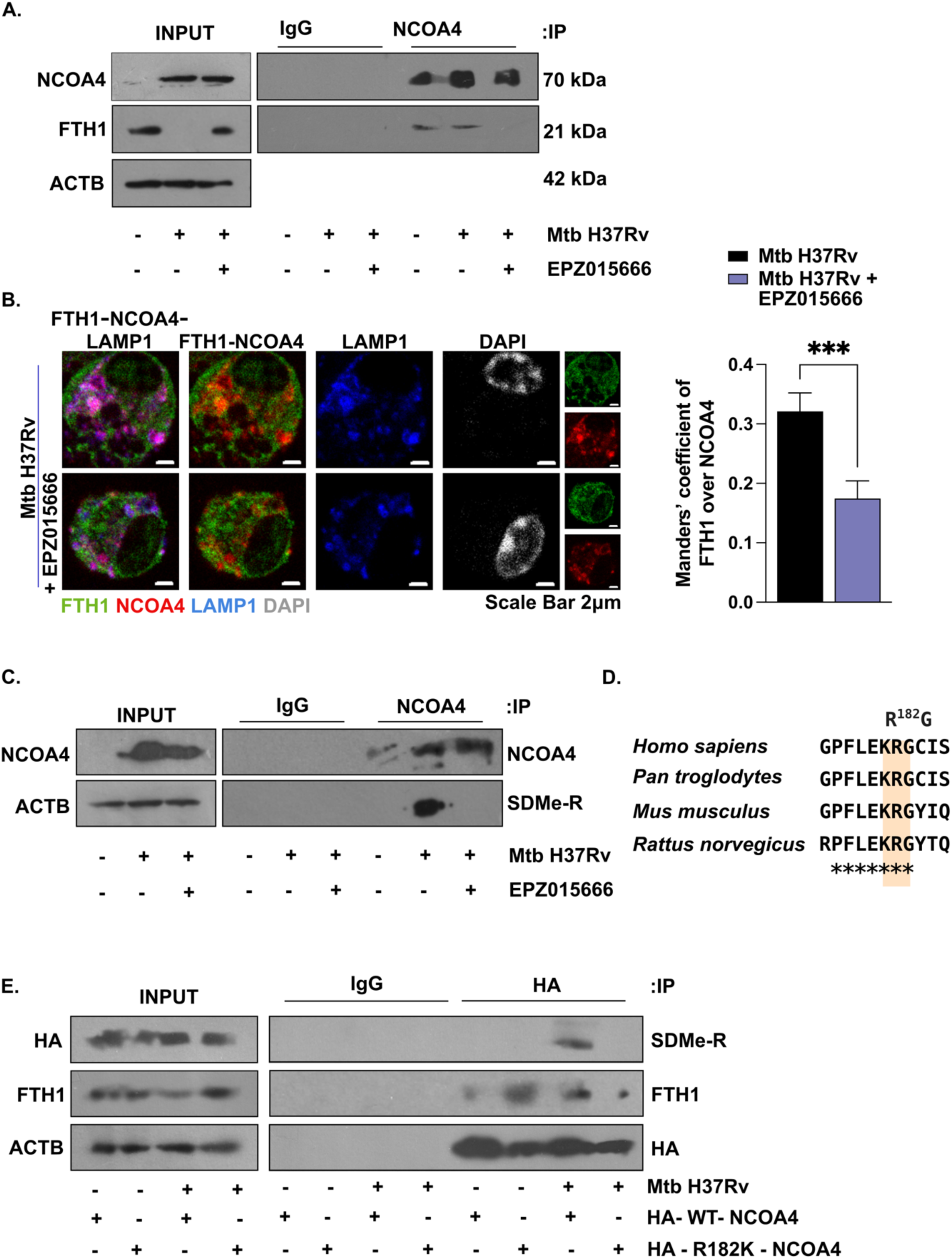
PRMT5 interacts with NCOA4 and methylates it. (**A**) Assessment of the interaction between NCOA4 and ferritin heavy chain by co-immunoprecipitation of whole cell lysates from murine macrophages pretreated with PRMT5 inhibitor (for 1 hour) at 24 hours post infection (**B**) Confocal images of murine macrophages pretreated with PRMT5 inhibitor (for 1 hour) stained for ferritin heavy chain (green), NCOA4 (red) lysosomes LAMP1 (blue) and nuclear staining (grey) at 24 hours post infection. Scale bars: 2 µm (left), and its quantification (right). (**C**) Assessment of extent of arginine methylation on NCOA4 by immunoprecipitation of whole cell lysates from murine macrophages pretreated with PRMT5 inhibitor (for 1 hour) at 24 hours post infection. (**D**) The putative arginine residue that is methylated in NCOA4 is conserved across different species. (**E**) Assessment of extent of arginine methylation on NCOA4 and its interaction with ferritin heavy chain by co-immunoprecipitation of whole cell lysates from RAW 264.7 cells transfected with NCOA4 OE vectors at 24 hours post infection. All immunoblotting is representative of three independent experiments. ACTB was used as a loading control. DAPI, 4’,6-diamidino-2-phenylindole; IP, immunoprecipitation; OE, overexpression. **, p<0.01; (Student’s t-test in B; GraphPad Prism 10.0). EPZ015666 (20 μM), PRMT5 inhibitor.

### Methylation of NCOA4 is important for Mtb-induced ferritinophagy and ferroptosis

We sought to then investigate whether PRMT5-dependent methylation on NCOA4 impacts ferroptosis. To this end, we found that the overexpression of R182K-NCOA4 could restore levels of ferritin heavy chain in Mtb infected cells when compared to cells overexpressing the NCOA4-WT (Figure 4A). Consistently, while the overexpression of NCOA4-WT could complement the intracellular iron pool in Mtb-infected cells, the R182K mutant NCOA4 could not, suggesting that the methylation of NCOA4 is crucial for ferroptosis-associated ferritinophagy (Figure 4B). Additionally, the overexpression of R182K-NCOA4 phenocopied PRMT5 inhibition and reduced levels of 4HNE in Mtb-infected macrophages (Figure 4C). This observation was further supported by reduced levels of Mtb in present in the culture supernatants of macrophages overexpressing the R182K-NCOA4 mutant indicating compromised Mtb release from infected cells (Figure 4D). Moreover, immunofluorescence studies confirmed that the ectopic expression of R182K-NCOA4 could reduce the colocalization of ferritin and NCOA4 with LAMP1 positive lysosomes (Figure 4E, S4H). Together, this highlights a critical role for R182 methylation of NCOA4 in Mtb-induced ferroptosis.

**Figure 4:**
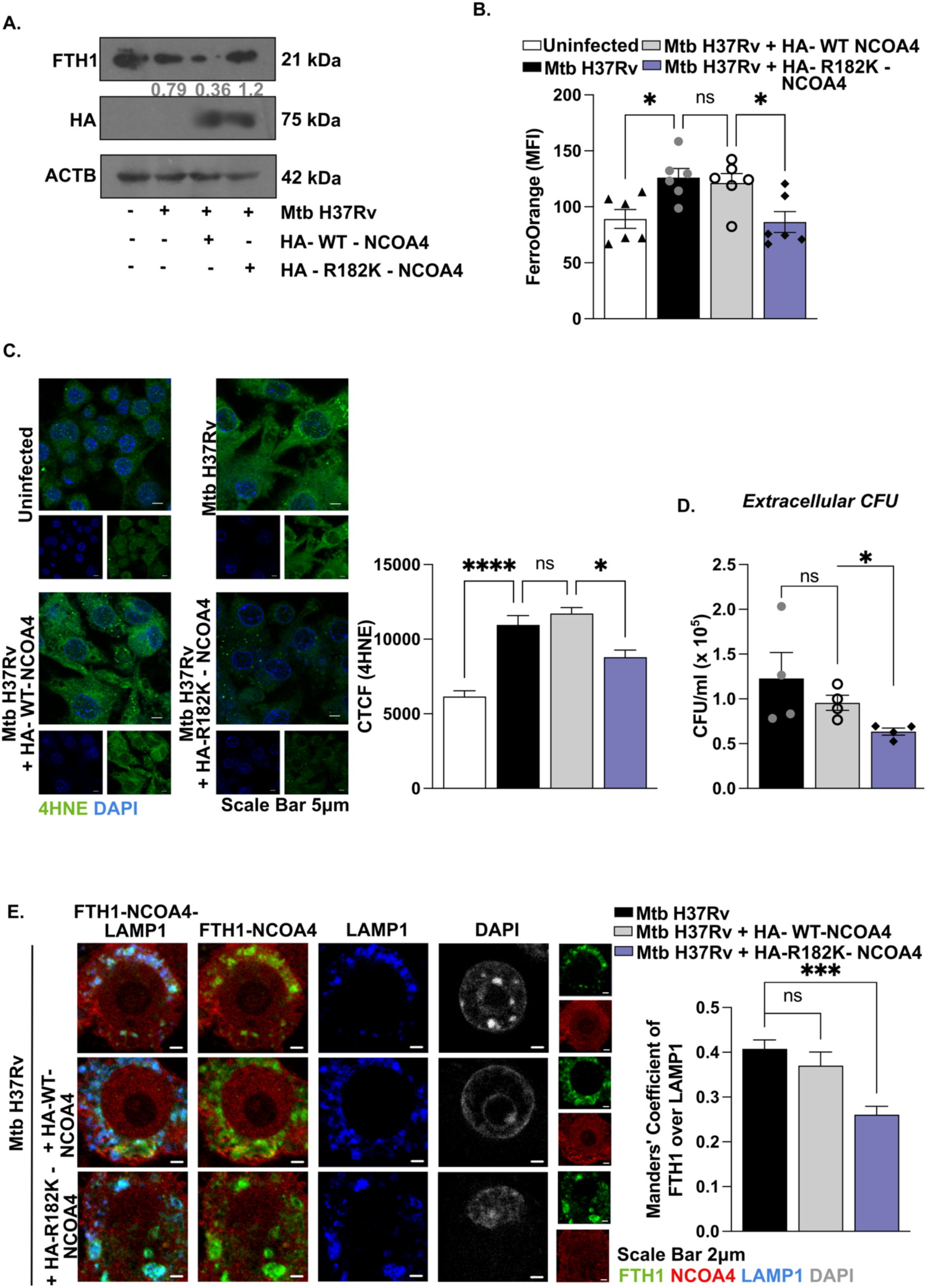
PRMT5-dependent methylation of NCOA4 is indispensable for ferritinophagy and subsequent ferroptosis during Mtb infection. (**A-G**) RAW 264.7 cells were transfected with NCOA4 OE vectors. (**A**) Assessment of the levels of ferritin heavy chain at 24 hours post infection by immunoblotting. (**B**) Estimation of intracellular labile ferrous iron at 24 hours post infection. (**C**) Confocal images stained for 4HNE (green) and nuclear staining (blue) at 24 hours post infection. Scale bars: 5 µm (left), and its quantification (right). (**D**) Examination of the release of live mycobacteria from necrotic cells by CFU quantification in culture supernatants at 4 days post infection. (**D**) Confocal images stained for ferritin heavy chain (green), NCOA4 (red) lysosomes LAMP1 (blue) and nuclear staining (grey) at 24 hours post infection. Scale bars: 2 µm (left), and its quantification (right). All immunoblotting and immunofluorescence data are representative of three independent experiments. ACTB was used as a loading control. MFI, mean fluorescence intensity; OE, over expression; DAPI, 4’,6-diamidino-2-phenylindole; CTCF, corrected total cell fluorescence; CFU, colony forming units. *, p< 0.05; **, p<0.01; ***, p<0.001; ****, p < 0.0001 (One-way ANOVA in B, C, E; Student’s t-test in D; GraphPad Prism 10.0). Below each panel, the quantification of the blots normalized to the loading control has been indicated.

### Methylation of NCOA4 affects its nuclear localization and function as a co-regulator of nuclear receptors

Apart from its role in iron homeostasis, NCOA4 was initially identified to interact with the androgen receptor (AR) and serve as a coactivator capable of enhancing AR-mediated transcription(*44*). Since, studies have identified that NCOA4 interacts with several other nuclear receptors including vitamin D3 receptor, estrogen receptor, and peroxisome proliferator-activated receptor (PPAR)αγ(*45*, *46*). Consistently, a previous investigation has identified that the cytoplasmic retention of NCOA4 may undermine the ligand-dependent effector functions of these nuclear receptors(*31*). Interestingly, the site of arginine methylation in mouse and its homologous residue in human NCOA4 lies between the two ARA70 binding domains that are essential for its interaction with nuclear receptors (Figure S5A). Thus, to understand the comprehensive function of NCOA4 methylation, we aimed to investigate whether it affects its nuclear localization. To verify this, we analyzed levels of NCOA4 in the nucleus and cytoplasm of macrophages infected with Mtb using immunofluorescence. We observed that the nuclear localization of NCOA4 is compromised upon Mtb infection (Figure S5B, C). However, this effect was reversed upon PRMT5 inhibition, indicating that PRMT5-mediated methylation of NCOA4 might result in the cytoplasmic retention of NCOA4 and thereby disrupt its coactivator function (Figure S5D, E). Consistent with this, the proportion of NCOA4 in the nucleus of Mtb-infected cells was also reversed upon the overexpression of R182K-NCOA4 (Figure 5A, B). Motivated by these observations, we assessed the effect of NCOA4 methylation on its interaction and downstream activity as a co-activating receptor. In this regard, we specifically assessed for its interaction with the vitamin D3 receptor, an association that has been previously demonstrated to enhance the transcriptional response of the 1,25-dihydroxyvitamin D₃ (1,25(OH)₂D₃)-induced signaling(*47*). Supplementation of vitamin D3 has also been reported to augment the antimycobacterial functions of Mtb-infected macrophages (*48*, *49*). Importantly, PRMT5 has also been identified to repress 1,25(OH)₂D₃-induced gene expression(*50*). In line with this, we could observe a marked decrease in the association between NCOA4 and vitamin D3 receptor post Mtb infection and 1,25(OH)₂D₃ supplementation (Figure S6A). Further, consistent with our earlier findings, the inhibition of PRMT5 could rescue their interaction (Figure 5C). Further, we could observe a marked increase in the 1,25(OH)₂D₃-induced nuclear localization of NCOA4 upon PRMT5 inhibition as opposed to the Mtb-infected cells (Figure 5D, S6B). Together, these findings suggest that PRMT5-dependent NCOA4 methylation could affect its nucleo-cytoplasmic localization and activity as a co-activating receptor.

**Figure 5:**
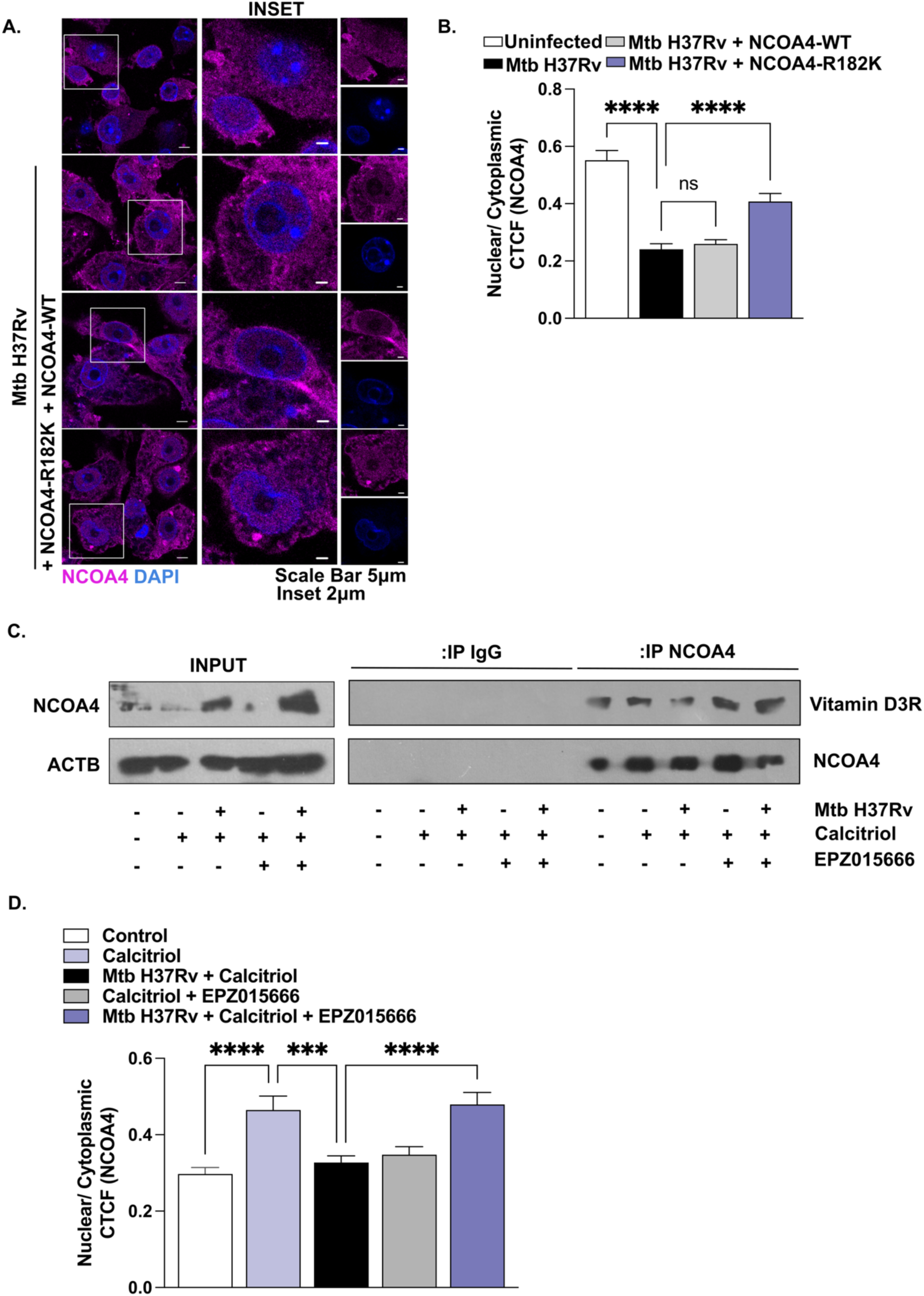
NCOA4 R182 methylation promotes its cytoplasmic retention and compromises its interaction with vitamin D3 receptor. (**A, B**) RAW 264.7 cells were transfected with NCOA4 OE vectors. Confocal images stained for NCOA4 (magenta) and nuclear staining (blue) at 24 hours post infection (**A**) and the quantification of the nuclear cytoplasmic localisation (**B**). (**C**) Assessment of the interaction between NCOA4 and vitamin D3 receptor by co-immunoprecipitation of whole cell lysates from murine macrophages pretreated with Calcitriol and PRMT5 inhibitor (for 1 hour) at 24 hours post infection. (D) Quantification of the nuclear cytoplasmic localisation of NCOA4 in murine macrophages pretreated with Calcitriol and PRMT5 inhibitor (for 1 hour) at 24 hours post infection by immunostaining. All immunofluorescence and immunoblotting data are representative of three independent experiments. DAPI, 4’,6-diamidino-2-phenylindole; CTCF, corrected total cell fluorescence; EPZ015666 (20 μM), PRMT5 inhibitor; Calcitriol (10nM), 1,25(OH)₂D₃. ***, p<0.001; ****, p<0.0001 (One-way ANOVA in B, D; GraphPad Prism 10.0).

### PRMT5 inhibition alongside vitamin D3 supplementation therapy helps combat tuberculosis and serve as potential adjunct therapies

Next, to understand the implications of NCOA4-mediated vitamin D3 signalling in clearance of Mtb, we assessed for the status of the downstream consequences of 1,25(OH)_2_D_3_ supplementation upon conditions of PRMT5 inhibition. Specifically, as vitamin D3 supplementation has been identified to overcome the early endosome-like arrest of Mtb and deliver it to the autolysosomes(*49*), we initially assessed for the status of autophagy in these cells. Consistently, PRMT5 inhibition could significantly enhance the 1,25(OH)₂D₃-induced colocalization of Mtb with lysosomes (Figure 6A, B). To understand the relevance of our observations in the treatment of tuberculosis, we utilized an animal model wherein BALB/c mice were infected with Mtb H37Rv and administered with 1,25(OH)₂D₃ alone or a combination of PRMT5 inhibitor and 1,25(OH)₂D₃ as indicated (Figure 6C). Interestingly, the combinatorial drug regime with vitamin D3 alongside PRMT5 inhibition could enhance the beneficial effects of 1,25(OH)₂D₃ supplementation and could effectively clear Mtb from the lungs, and spleen of infected mice (Figure 6D, E). Concomitantly, the combinatorial regime could also improve TB pathology and reduce the formation of hallmark TB granuloma-like lesions within the lungs as indicated by the blinded assessment of H and E-stained lung sections and assessment of granuloma-like fraction within the infected lungs (Figure 6F).

**Figure 6:**
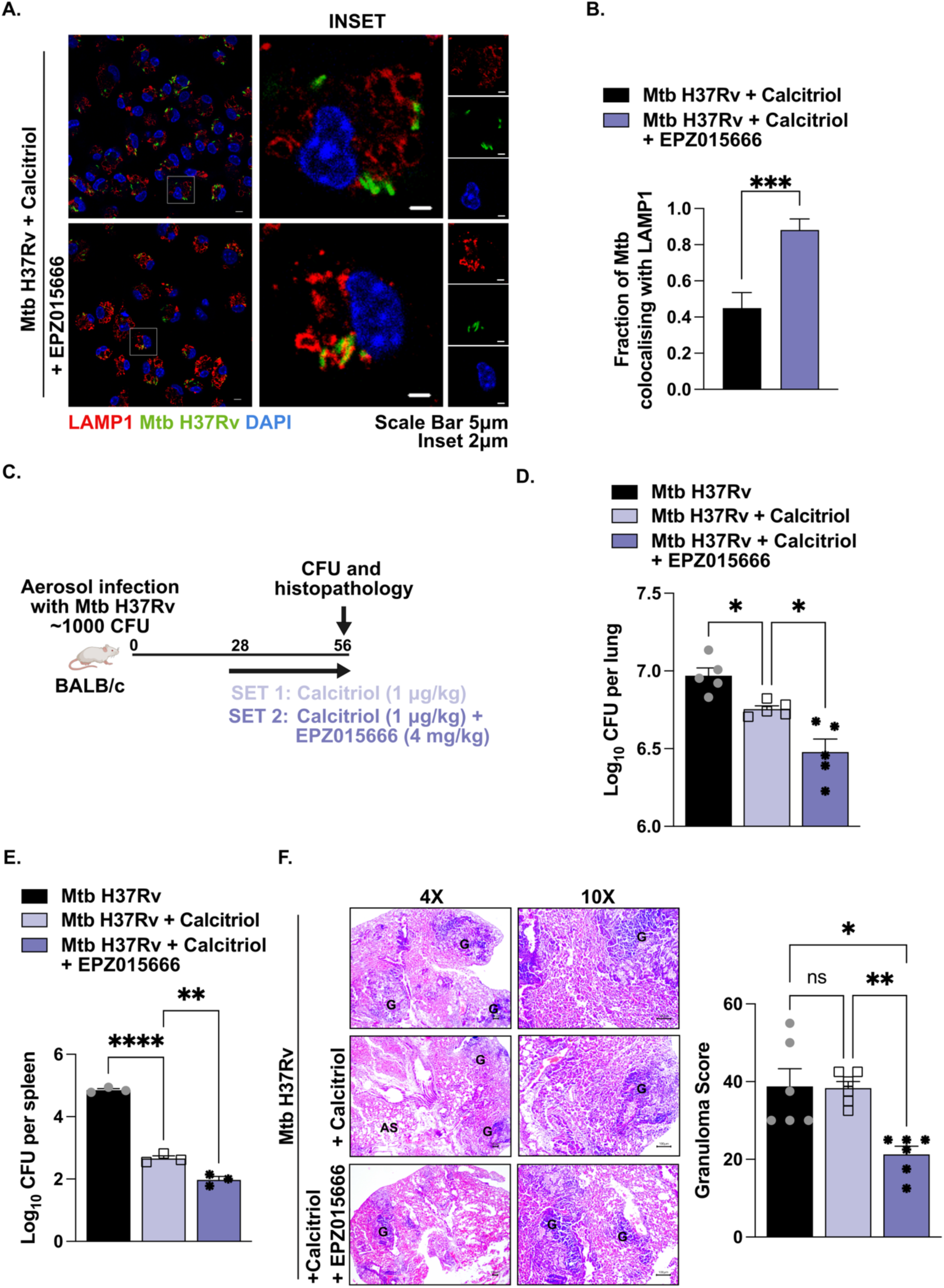
PRMT5 inhibition enhances vitamin D3-mediated immunity against Mtb. (**A, B**) Confocal images stained for lysosomes LAMP1 (red) and nuclear staining (blue) infected with Mtb H37Rv-GFP (green) in murine macrophages pretreated with PRMT5 inhibitor (for 1 hour) at 24 hours post infection (**A**) and its respective quantification (**B**). (**C-F**) BALB/c mice were infected with Mtb H37Rv and treated with Calcitriol (1μg/kg) q.d and PRMT5 inhibitor EPZ015666 (4mg/kg) q.o.d. as indicated in (**C**) (Number of mice per group = 5). (**D, E**) Enumeration of mycobacterial burden in the lungs (**D**) and spleen (**E**) of infected and inhibitor treated mice by plating organ homogenates on 7H11 plates. (**F**) Analysis of TB pathology (granulomatous lesions) in the lung sections with H and E staining (left) and its quantification by a pathologist in a blinded manner (right). Specific regions of the H and E-stained sections were analysed by the pathologist for evaluation of the granuloma fraction. Accordingly, the portions have been demarcated in the images; Magnification, 4X (left panel); 10X (right panel); Scale bar for H and E images, 100μm; G, granuloma; AS, alveolar space; DAPI, 4’,6-diamidino-2-phenylindole; CFU, colony forming units; q.o.d., *quaque altera die*, every other day; q.d., *quaque die*, once a day. EPZ015666 (20 μM), PRMT5 inhibitor; Calcitriol (10nM), 1,25(OH)₂D₃. *, p<0.05; ***, p<0.001 (One-way ANOVA in E; Student’s t-test in C; GraphPad Prism 10.0).

Thus, we report the PRMT5-dependent arginine methylation of the nuclear co-activating receptor, NCOA4, during Mtb infection. This post-translational modification contributed to the induction of ferroptosis and repression of vitamin D3 signalling during TB, thereby aiding in mycobacterial survival, both in vitro and in vivo. Together, our experimental evidence highlights the key role of PRMT5-NCOA4 in regulating the pathogenesis of TB.

## Discussion

Mtb infection is a leading cause of morbidity and annually accounts for nearly 1.4 million deaths worldwide. The success of Mtb is widely attributed to its ability of circumventing the host immune system and induction of cellular damage to facilitate its pathogenesis. In this regard, host-directed therapeutics (HDTs), aiming to exploit such intricate interactions between Mtb and the host, have garnered attention and appear promising in overcoming the unmet needs of conventional TB therapy(*51*). Specifically, HDTs as adjunct therapy can augment the host immune response, reduce exacerbated inflammation and tissue damage, thereby improving and accelerating TB treatment. To this end, ferroptosis has been proposed as a potential target for HDT against TB, given its crucial role in Mtb-induced tissue damage and dissemination. During Mtb infection, ferroptosis is driven by modulated levels of BACH1 (BTB domain and CNC homologue 1), a suppressor of NRF2 (Nuclear factor erythroid 2–related factor 2), and GPX4, resulting in impaired antioxidant responses (*23*). A host of other studies have substantiated these findings and identified bacterial effector proteins, including PTPA (protein tyrosine phosphatase A) and Rv1324, that alter host pathways and induce ferroptosis in Mtb-infected cells (*25*, *52*). Despite these advances, significant gaps remain in our understanding of Mtb-induced ferroptosis.

Our study provides the first evidence for NCOA4-driven ferritinophagy in inducing ferroptosis during TB and identifies key regulators of the process. Importantly, we identify a novel post-translational modification on NCOA4, as an indispensable player in the induction of ferritinophagy during Mtb infection. This arginine methylation was conferred by PRMT5 and the overexpression of the methylation deficient R182K NCOA4 mutant could drastically reduce the NCOA4 ferritin interaction and rescue cells from iron overload. NCOA4 structure is not well defined apart from the N-terminal coil-coil domain proposed to play a role in protein interactions. While the R182 residue on the mouse protein and its homologous residue in human NCOA4 is present within the disordered region adjacent to the N-terminal domain, the mechanism by which it regulates the molecular interaction between NCOA4, and ferritin needs further investigation.

Protein arginine methylation catalysed by various members of the PRMT family are being increasingly reported to regulate ferroptosis. For instance, type I PRMTs that confer an asymmetric di methylation on arginine such as PRMT1 has been reported to regulate diverse aspects of lipid metabolism, iron homeostasis and antioxidant responses (*35*, *53*, *54*) The type III PRMT7, that catalyses the transfer of a mono methylation on arginine residues has also been reported to regulate levels of ACSL4 and contribute to ferroptosis(*55*). In this study, we provide the first evidence of symmetric demethylation of arginine catalysed by the type II PRMT, PRMT5, in regulating ferritinophagy. Interestingly, we observe that the methylation could overcome the homeostatic control of iron levels and induce ferritin degradation despite the status of iron availability. While PRMT5 is the major enzyme catalysing the symmetric di methylation of arginine in mammalian cells, the role of the other type II enzyme, PRMT9 has not been investigated. Similarly, molecular crosstalk between distinct PRMTs might also play a role and would require further investigation. In this regard, a study has reported the interaction between PRMT4 and NCOA4 in acute kidney injury (*56*) While PRMT4 does not methylate NCOA4, it might be interesting to evaluate whether it synergizes with PRMT5 in regulating ferritinophagy.

Ferritinophagy is conventionally known to be dependent on NCOA4 levels which are in turn tightly regulated by the cellular iron levels ensuring that the induction of ferritinophagy depends on iron availability and demand (*57*). This is facilitated by the iron-induced interaction between HERC2 and NCOA4 and the concomitant proteasomal degradation of the cargo receptor(*58*). However, in conditions such as Mtb infection, wherein this regulatory machinery is compromised, NCOA4 is stable despite iron availability and is aberrantly accumulated within the cell(*30*). In such conditions, other cellular mechanisms must exist to prevent iron-induced toxicity and might be effectuated by the modulation of distinct post-translational modifications. Previous research has shown that ATM-dependent phosphorylation or interaction with STING facilitates the interaction between NCOA4 and ferritin and contributes to the iron availability in the cell(*31*, *32*). While our study demonstrates a key role for PRMT5 in inducing ferroptosis in the iron replete conditions upon Mtb infection, further studies will also help to understand whether PRMT5-dependent methylation of NCOA4 is a general machinery regulating ferritinophagy during iron starvation.

Additionally, NCOA4 can also serve as the co-activating factor for other steroid receptors including the androgen receptor (AR), aryl-hydrocarbon receptor (AhR), and peroxisome proliferator-activated receptor (PPAR) which are also well implicated during Mtb infection. Our studies demonstrate that the methylation of NCOA4 can promote its cytoplasmic retention and impair its activity in the vitamin D3-induced immune response. Importantly, the inhibition of PRMT5 could reverse this effect and improve vitamin D3-induced immunity against TB. This opens up new prospects of study as utilization of vitamin D3 supplementation therapy has been widely proposed as a HDT against TB. However, the clinical findings regarding its protective role during TB remains inconsistent and our understanding of the process remains incomplete. Our results thus identify a Mtb-induced suppression of vitamin D3 signalling and report that the inhibition of PRMT5 alongside the supplementation of vitamin D3 could enhance the effect of vitamin D3 and thereby serve as novel and promising target in therapy.

In summary, our study identified a previously undiscovered function of PRMT5 in regulating the interaction between NCOA4 and ferritin. The methylation of the arginine residue on NCOA4 induced ferritinophagy and ferroptosis in Mtb infected cells and is pathologically important in TB. Furthermore, we find a significant reduction in Mtb burden and TB pathogenesis upon administration of PRMT5 inhibitor alongside vitamin D3 supplementation. Given the importance of host iron status and vitamin D3 supplementation that has been long known to affect Mtb survival, we believe that strategies aimed at targeting these molecular mediators will serve as a promising host-directed therapy against TB.

## Materials and Methods

### Ethics statement

All experiments involving mice were carried out after obtaining approval from Institutional Ethics Committee for animal experimentation, Indian Institute of Science (IISc), Bengaluru, India (Approval Number: CAF/ETHICS/062/2024). The animal care and use protocol adhered were approved by national guidelines of the Committee for Control and Supervision of Experiments on Animals (CCSEA), Government of India. Experiments with virulent mycobacteria (Mtb H37Rv) were approved by the Institutional Biosafety Committee, Indian Institute of Science (IISc), Bengaluru, India (Approval Number: IBSC/IISC/KNB /18/2023-24).

### Cells and mice

Four to six weeks old, male, and female BALB/c mice were utilized for all experiments. Mice were procured from The Jacksons Laboratory and maintained at the Central Animal Facility (CAF) in Indian Institute of Science (IISc) under 12-hour light and dark cycle. For in vitro experiments, mouse peritoneal macrophages were utilized. Briefly mice were injected with 8% Brewer’s thioglycollate (HiMedia Laboratories, M019), and peritoneal exudates were isolated in ice-cold PBS after four days. The cells were seeded in tissue culture dishes and adherent cells were utilized as peritoneal macrophages. RAW 264.7 mouse monocyte-like cell line was obtained from American Type Culture Collection (ATCC). Primary macrophages and RAW 264.7 macrophage cell line were cultured in Dulbecco’s Modified Eagle Medium (DMEM, Gibco, Thermo Fisher Scientific, 12100061) supplemented with 10% heat inactivated Fetal Bovine Serum (FBS, Gibco, Thermo Fisher Scientific, 10270106) and maintained at 37°C in 5% CO2 incubator. THP-1 human monocyte cell line line was obtained from American Type Culture Collection (ATCC). THP-1 cells were cultured in RPMI-1640 synthetic medium (Gibco, Thermo Fisher Scientific, 31800022) with 10% heat inactivated FBS supplement. Monocytes were differentiated by treatment with 20 ng/mL phorbol 12-myristate 13-acetate (PMA; Sigma-Aldrich, P1585) for 18 h. Thereafter, cells were rested for two days to ensure their reversion to a resting phenotype before infection.

### Bacteria

Virulent strain of Mtb (Mtb H37Rv) was a kind research gift from Prof. Amit Singh, Department of Microbiology and Cell Biology, and Centre for Infectious Disease Research, IISc. Mycobacteria were cultured in Middlebrook 7H9 medium (BD Difco, 271310) supplemented with 10% OADC (oleic acid, albumin, dextrose, catalase). Single-cell suspensions of mycobacteria were obtained by passing mid log phase culture through 30-gauge needle 5 times each and used for infecting cells at multiplicity of infection 10. The studies involving virulent mycobacterial strains were carried out at the biosafety level 3 (BSL-3) facility at Centre for Infectious Disease Research (CIDR), IISc.

### Reagents and antibodies

All general chemicals and reagents were procured from Sigma-Aldrich/ Merck Millipore, HiMedia and Promega. Tissue culture plastic ware was purchased from Nunc Cell Culture, Thermo Fisher Scientific and Tarsons India Pvt. Ltd. siRNAs were obtained from Eurogentec against *Ncoa4*, *Prmt5*. 4′,6-Diamidino-2-phenylindole dihydrochloride (DAPI) were procured from Invitrogen, Thermo Fisher Scientific, D1306. HRP-tagged anti-β-ACTIN (A3854) and anti-MAP1LC3B (L7543) was procured from Sigma Aldrich. Anti-FTH1 (3998), anti-PRMT5 (2252), anti-SdMe-R (13222), anti-VitD3R (12550), anti-LC3A/ B (4108), anti-SQSTM1 (5114) antibodies were procured from Cell Signaling Technology (USA). Anti-NCOA4 (ab86707), anti-4HNE (ab46545), antibody was sourced from Abcam. Anti-HA (26183) and anti-LAMP1 (14-1071-82) antibody was obtained from Invitrogen, Thermo Fisher Scientific. Anti-GPX4 (NBP2-76933) antibody was procured from Novus Biologicals. Anti-FTH1 (MAD021Mu21) antibody was obtained from Cloud Clone HRP conjugated anti-rabbit IgG/anti-mouse IgG was obtained from Jackson ImmunoResearch (USA).

### Treatment with pharmacological reagents

Macrophages were pre-treated with the following reagents one hour prior to infection with Mtb H37Rv: EPZ015666 (Tocris, 6516; Cayman Chemicals, 17285; 20 μM); Calcitriol (Cayman Chemicals, 71820; 10nM); Chloroquine (Sigma-Aldrich, C6628; 10 μM); MG132 (Sigma-Aldrich, 10 μM).

### Estimation of Intracellular labile iron pool

Labile intracellular iron was measured by using the BioTracker FerroOrange Live Cell Dye (EMD Millipore Corp. SCT210). Cells post treatment and/or infection were washed with 1X PBS. FerroOrange dye at a concentration of 1μM was added and incubated at 37°C for 30 minutes in the CO2 incubator. Cells were washed with 1X PBS and the FerroOrange florescence was analyzed with a fluorescence microplate reader (SpectraMax M3 at Ex/Em = 542 nm/ 572 nm).

### Immunofluorescence

Cells were fixed with 3.7% formaldehyde for 30 min. at room temperature. Fixed samples were blocked with 2% BSA in PBST (containing 0.02% saponin) for 1 h. Microtome sections (5 μm) were obtained from paraffin-embedded mouse lung tissue samples. Deparaffinized sections were subjected to antigen retrieval by boiling in 10 mM citrate buffer (pH 6.0) for 15 min; treated with 1% H2O2 for 10 min in dark and blocked with 2% BSA in PBST for 1 h at room temperature. After blocking, samples were stained with the indicated primary antibodies in 2% BSA in PBST at 4°C overnight. After incubation, the cells/ sections were washed with PBST followed by incubation with Alexa Fluor 488- (Invitrogen, A27034), Alexa Fluor 555- (CST, 4413), CF 555 (Sigma Aldrich, SAB4600070) or Alexa Fluor 647- (Invitrogen, A32733; Jackson ImmunoResearch, 115-605-003) conjugated secondary antibodies for 2 h and nuclei were stained with DAPI (Invitrogen, D1306). The samples were mounted on glycerol. Confocal images were taken with Zeiss LSM 710 Meta confocal laser scanning microscope (Carl Zeiss AG, Germany) using a plan-Apochromat 63X/1.4 Oil DIC objective (Carl Zeiss AG, Germany). For quantitative estimation of the results, at least 100 cells from different fields were analyzed. For the Mean Fluorescence Intensity (MFI) analysis, ImageJ Fiji was utilized to calculate the maximum intensity projections of the Z-stacks. Using free hand selection tool, cells were selected to measure the area-integrated intensity and mean grey value. The area around the cells without fluorescence was used to calculate the background values. Corrected Total Cell Fluorescence (CTCF) was calculated using the following formula: CTCF = Integrated intensity - (area of selected cell X Mean florescence of background reading). Manders’ correlation coefficient between two channels was quantified using JACoP plugin of ImageJ Fiji software.

### CellROX staining for determining oxidative stress

CellROX Deep Red Reagent (Invitrogen, C10422) was used to determine oxidative stress in macrophages as per manufacturer’s instructions. Briefly, macrophages were treated with CellROX Deep Red Reagent at a final concentration of 5 μM, diluted in DMEM without phenol red. The treated cells were incubated for 30 min at 37 °C in the CO2 incubator. Cells were then washed with PBS thrice, followed by fixation with 3.7 % formaldehyde for 15 min. Nuclei were stained with DAPI and images were captured in Zeiss LSM 710 confocal laser scanning microscope (Carl Zeiss AG, Germany) using a plan-Apochromat 63X/1.4 Oil DIC objective (Carl Zeiss AG, Germany). The quantitative estimation of the results was performed as described previously.

### Lipid staining for estimation of lipid peroxidation

Lipid peroxidation was assessed using BODIPY 493/503 (Invitrogen, D3922) and BODIPY 665/676 (Invitrogen, B3932). The cells were fixed with 3.7% formaldehyde for 30 min. After 3 washes with PBS, the cells were stained with 10 μg/mL BODIPY 493/503 and BODIPY 665/676 for 30 min in the dark. Subsequently, the cells were washed, and nuclei were stained with DAPI. After washing with PBS, samples were mounted on glycerol and visualized and analyzed by confocal microscopy. Confocal images were taken with Zeiss LSM 710 Meta confocal laser scanning microscope (Carl Zeiss AG, Germany) using a plan-Apochromat 63X/1.4 Oil DIC objective (Carl Zeiss AG, Germany). The quantitative estimation of the results was performed as described previously. The ratio of unoxidized lipids to neutral lipids was calculated as CTCF (BODIPY 665/676)/ CTCF (BODIPY 493/503) for every cell.

### Measurement of cell death

Necrotic cell death was assessed by measuring the release of LDH in the supernatants from macrophage cultures using CyQUANT LDH Cytotoxicity Assay (Invitrogen, C20300) according to the manufacturer’s instructions. The absorbance at 490nm was measured using a microplate reader (SpectraMax M3). The percent LDH released was calculated using the following formula: [Absorbance of sample/ Absorbance of lysed replicate (maximum LDH)] * 100

### Bacterial enumeration

Cells were infected with Mtb H37Rv at MOI 10 for 4h. After 4h of infection, macrophages were washed thoroughly with PBS to eliminate the presence of the extracellular bacterium. Cells were then supplemented with a medium containing amikacin (HiMedia, CMS644) at 0.2mg/ml for 2h to deplete any extracellular bacterium. Cells were washed with PBS and supplemented with DMEM medium at this time, taken as 0h. Duplicates were maintained with respective inhibitors in the antibiotic-free medium. Extracellular CFU counts were determined at the indicated time points. Extracellular bacteria were evaluated by plating serial dilutions of culture supernatants onto Middlebrook 7H11 (BD Difco, 283810) agar plates supplemented with 0.5% (vol/vol) glycerol and 10% (vol/vol) OADC enrichment media. Colonies were counted after 21 d of incubation of 7H11 agar plates at 37°C.

### in vivo mouse model for TB and treatment with pharmacological inhibitor

BALB/c mice (n = 30) were infected with mid-log phase Mtb H37Rv, using a Madison chamber aerosol generation instrument calibrated to 1000 CFU/animal. Aerosolized animals were maintained in securely commissioned BSL3 facility. Post 28 days of established infection, mice were administered intra-peritoneal doses of EPZ015666 (4mg/kg) every alternate day over 28 days. On 56th day post inhibitor treatment, mice were sacrificed, the left lung lobe was homogenized in sterile PBS, serially diluted, and plated on 7H11 agar containing OADC to quantify CFU. Upper right lung lobes were fixed in formalin, and processed for hematoxylin and eosin staining, or immunofluorescence analyses. Also, specific lobes from the lungs of mice were homogenized for the extraction of RNA and protein.

### Hematoxylin and Eosin staining

Tissues fixed in formalin were handed over to consultant pathologist for blinded analyses. Briefly, microtome sections (5 μm) were obtained from paraffin-embedded mouse lung tissue samples. Deparaffinized and rehydrated sections were subjected to Hematoxylin staining followed by Eosin staining as per manufacturer instructions. After dehydrating, sections were mounted using permount. The pathologists have performed the analysis in a blinded manner based on the article by Palanisamy et al. Tuberculosis (Edinb). 2008.

### Lipid peroxidation measurement in the lungs of infected mice

Lipid peroxidation in lungs was assessed by using the TBARS assay kit (Cayman Chemicals, 703002) according to the manufacturer’s instructions. Briefly, mouse lung tissues were collected and homogenized in RIPA buffer. After centrifugation, SDS solution and the colour reagent was added to the supernatant from each sample. This mixture was then incubated at 95° for 1h, cooled to room temperature, centrifuged and added to 96-well plate to calculate the absorbance at 532 nm in a microplate reader (SpectraMax M3). The slope of the standard curve was used to calculate the amount of MDA (μM) in the samples.

### Immunoblotting

Cells post treatment and/or infection were washed with 1X PBS. Whole cell lysate was prepared by lysing in RIPA buffer [50 mM Tris-HCl (pH 7.4), 1% NP-40, 0.25% sodium deoxycholate, 150 mM NaCl, 1 mM EDTA, 1 mM PMSF, 1 μg/mL each of aprotinin, leupeptin, pepstatin, 1 mM Na3VO4, 1 mM NaF] on ice for 30min. Total protein from whole cell lysates was estimated by Bradford reagent (Sigma Aldrich, B6916). Equal amount of protein was resolved on 12% SDS-PAGE and transferred onto PVDF membranes (Merck Millipore, IPVH00010) by semi-dry immunoblotting method (Bio-Rad). 5% non-fat dry milk powder in TBST [20 mM Tris-HCl (pH 7.4), 137 mM NaCl, and 0.1% Tween 20] was used for blocking nonspecific binding for 60 min. After washing with TBST, the blots were incubated overnight at 4°C with primary antibody diluted in TBST with 5% BSA. After washing with TBST, blots were incubated with anti-rabbit IgG secondary antibody conjugated to HRP antibody (Jackson ImmunoResearch, 111-035-045), or anti-mouse IgG secondary antibody conjugated to HRP antibody (Jackson ImmunoResearch, 115-035-003) for 6h at 4°C. The immunoblots were developed with enhanced chemiluminescence detection system (Perkin Elmer; BioRad, 1705061) as per manufacturer’s instructions. For developing more than one protein at a particular molecular weight range, the blots were stripped off the first antibody at 60°C for 5 min using stripping buffer (62.5 mM Tris-HCl, with 2% SDS 100 mM 2-Mercaptoethanol), washed with 1X TBST, blocked; followed by probing with the subsequent antibody following the described procedure. ACTB was used as loading control.

### Transient transfection studies

RAW 264.7 macrophages were transfected with the indicated constructs or mouse peritoneal macrophages were transfected with 100 nM each of non-targeting siRNA or specific siRNAs with the help of Lipofectamine 3000 (Invitrogen, L3000008) for 6h: followed by 24h recovery. Transfected cells were subjected to the required infections/ treatments for the indicated time points and processed for analyses.

### Immunoprecipitation assay

Immunoprecipitation assays were carried out following a modified version of the protocol provided by Millipore, USA. Treated samples were washed in ice cold PBS and gently lysed in RIPA buffer. The cell lysates obtained were subjected to pre-clearing with BSA-blocked Protein A beads (Bangalore Genei, India) for 30 min at 4°C and slow rotation. The amount of protein in the supernatant was quantified and equal amount of protein was used for pull down from each treatment condition; using Protein A beads pre-conjugated with the antibody of interest or isotype control IgG antibody. After incubation of the whole cell lysates with the antibody-complexed beads for 6h at 4°C on slow rotation, the pellet containing the bead-bound immune complexes were washed with RIPA buffer twice. The complexes were eluted by boiling the beads in Laemmli buffer for 10 min. The bead free samples were resolved by SDS-PAGE and the target interacting partners were identified by immunoblotting. Clean-Blot™ IP Detection Reagent (21230) was obtained from Thermo Scientific.

### Plasmids and constructs

pcDNA 3.1-NCOA4-HA OE and pcDNA 3.1-NCOA4-R182K-HA OE was synthesized from SynBio Technologies.

### Statistical analysis

Levels of significance for comparison between samples were determined by the student’s t-test and one-way ANOVA followed by Tukey’s multiple-comparisons. The data in the graphs are expressed as the mean ± S.E.M for the values from at least 3 or more independent experiments and P values < 0.05 were defined as significant. GraphPad Prism software (10.0 versions, GraphPad Software, USA) was used for all the statistical analyses.

## Acknowledgments

We acknowledge the Central Animal Facility, IISc for maintaining and providing mice used in the study. We thank the BSL-3 facility and staff for helping us in our in vitro and in vivo experiments with Mtb H37Rv. We thank Prof. Subba Rao Gangi Setty for his insights and guidance. We are also grateful to Prof. Balaji K.N.’s research group, majorly Awantika Shah and Atheena Abhayakumar for their critical comments on the manuscript. We acknowledge the help of Mahima and Dr. Salik Borbora for their guidance and insights on the manuscript.

## Funding

This work was supported by the Department of Biotechnology, Government of India BT/PR47843/MED/29/1631/2023, BT/PR41341/MED/29/1535/2020, BT/PR27352/BRB/10/1639/2017, BT/PR13522/COE/34/27/2015 (KNB); Department of Science and Technology, Government of India EMR/2014/000875 (KNB); Anusandhan National Research Foundation J. C. Bose National Fellowship SB/S2/JCB-025/2016, JBR/2021/000011 (KNB); Anusandhan National Research Foundation CRG/2019/002062, IRPHA-IPA/2021/000180 (KNB). The authors thank the DST-Fund for Improvement of S&T Infrastructure, University Grants Commission (UGC) Centre for Advanced Study, and DBT-IISc Partnership Program (phase II at IISc, BT/PR27952/INF/22/212/2018) for funding and infrastructure support. KNB also acknowledges the funds received as a part of infrastructure support to IISc as a part of the Institute of Eminence (IOE) scheme of the Government of India (IE/REDA-23-1757). Fellowships were received from CSIR and The Prime Minister’s Research Fellows Scheme (to SS). The funders had no role in study design, data collection and analysis, decision to publish or preparation of the manuscript.

## Competing interests

Authors declare that they have no competing interests.

## Data and materials availability

All data are available in the main text or the supplementary materials.

## Notes

### Competing Interest Statement

The authors have declared no competing interest.

