## Supplemental Figures for "Infection with *Mycobacterium tuberculosis* orchestrates the PRMT5-dependent methylation of NCOA4 to govern host ferroptosis"

#### Figure S1.

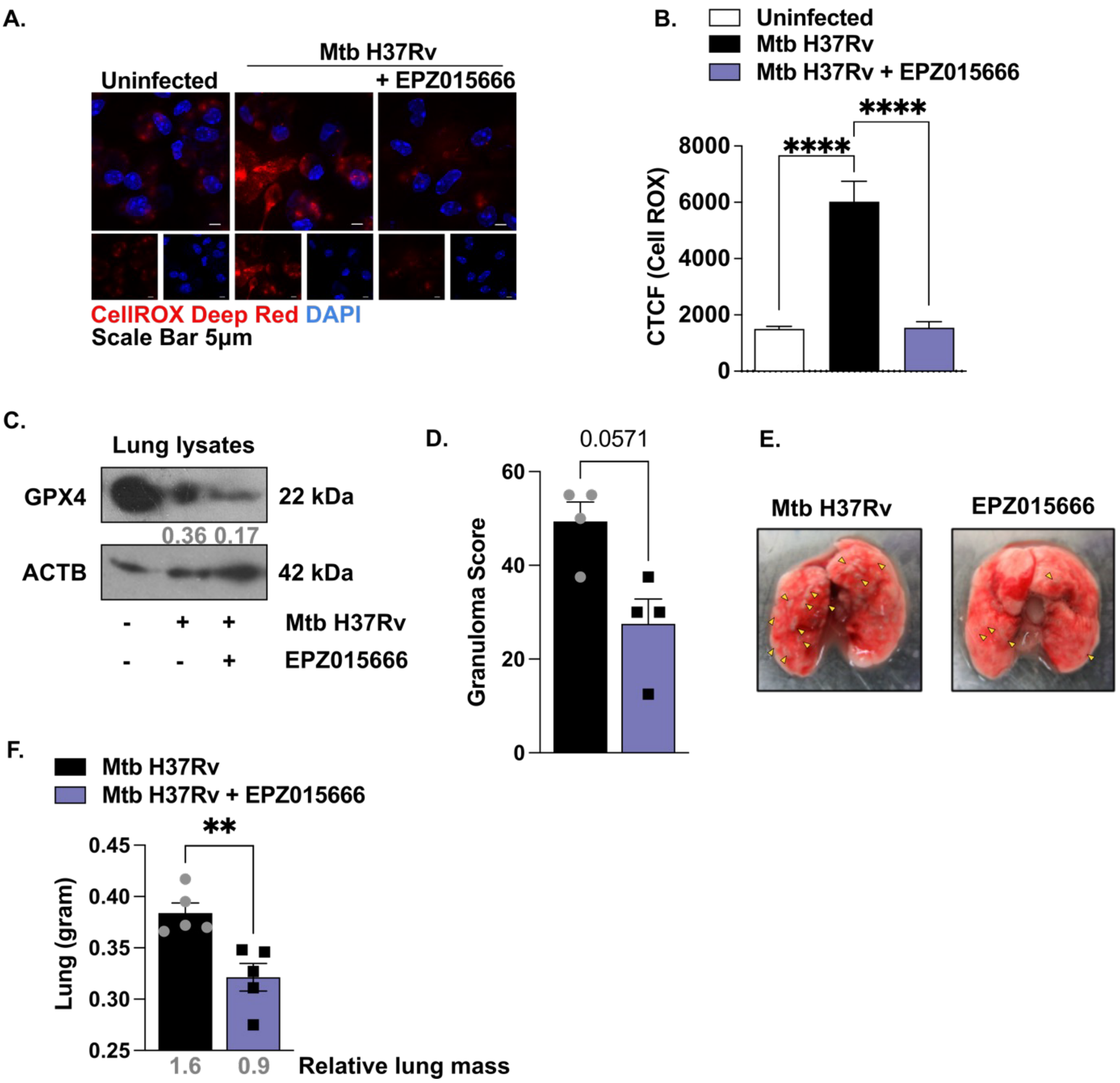


**Figure S1:** (**A-E**) Murine macrophages were treated with PRMT5 inhibitor for 1 hour, followed by infection with Mtb H37Rv for 24 hours. (**A, B**) Confocal image for the assessment of the intracellular oxidative stress using the CellROX Deep Red reagent (**A**) and its quantification (**B**). (**C-F**) BALB/c mice were infected with Mtb H37Rv and treated with PRMT5 inhibitor EPZ015666 (4mg/kg) q.o.d. as indicated in (**Figure 1D**) (Number of mice per group = 5) (**C**) Assessment of GPX4 levels by immunoblotting. (**D**) Analysis of the granuloma score in the lungs by H and E staining and quantification by a pathologist in a blinded manner. (**E**) Visualization of the gross pathology of the lungs from the images of whole lungs. Yellow arrows indicate visible granulomatous lesions (**F**) Lung weight of infected and PRMT5 inhibitor treated mice was plotted. The average relative wet mass of each lung is indicated below the bars. All immunofluorescence and immunoblotting data are representative of three independent experiments. ACTB was used as a loading control. DAPI, 4',6-diamidino-2-phenylindole; CTCF, corrected total cell fluorescence; EPZ015666 (20 µM), PRMT5 inhibitor. **, p<0.01; ****, p < 0.0001 (One-way

ANOVA in B; Student's t-test in D, F; GraphPad Prism 10.0). Below each panel, the quantification of the blots normalized to the loading control has been indicated.

**Figure S2.**

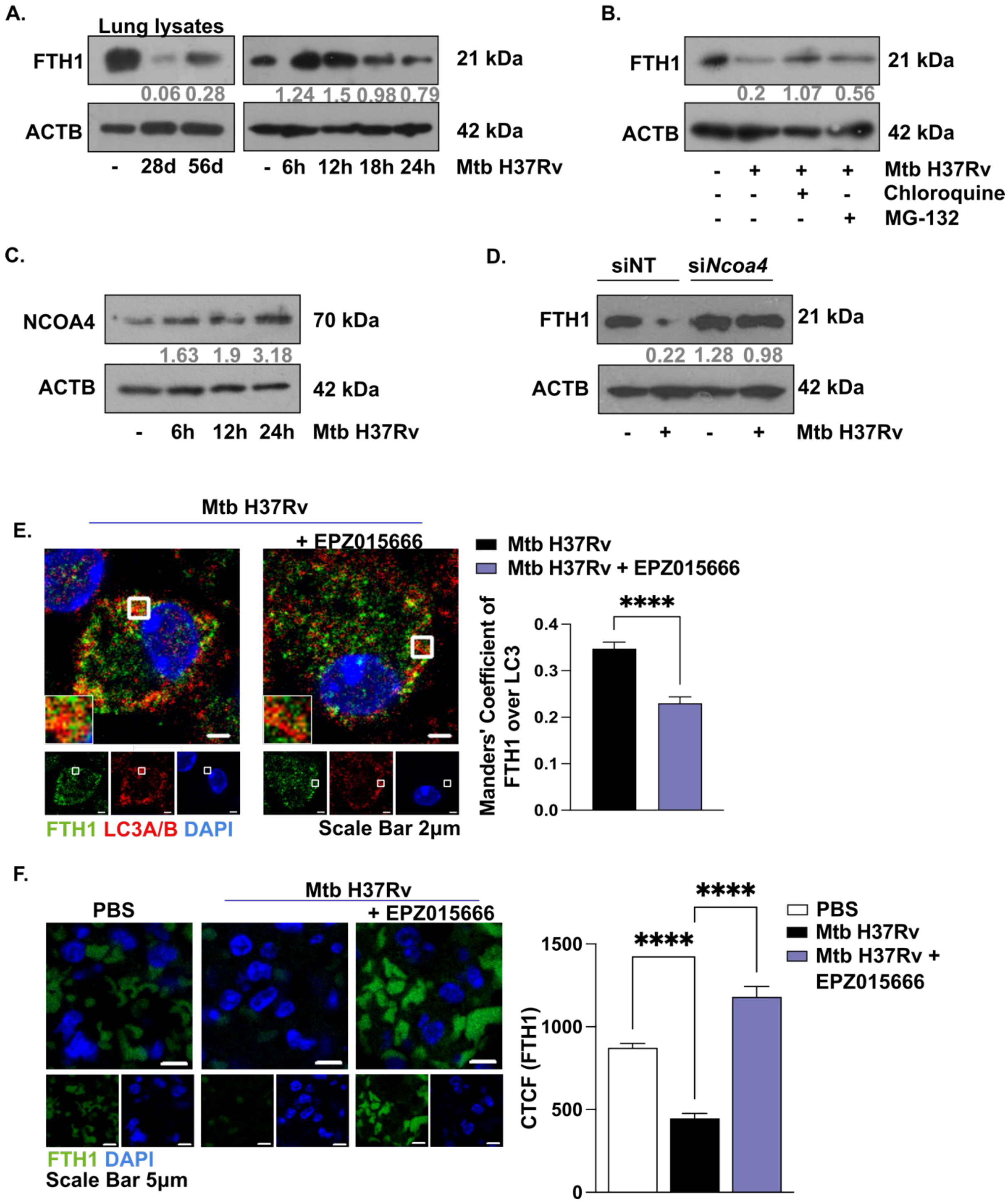


**Figure S2:** (**A**) Assessment of ferritin heavy chain levels in the lung homogenates of uninfected and infected BALB/c mice (left) and in murine macrophages infected with Mtb H37Rv for the indicated time points (right) by immunoblotting. (**B**) Assessment of ferritin heavy chain levels in murine macrophages pretreated with

Chloroquine (autophagy inhibitor) and MG-132 (proteasomal inhibitor) (for 1 hour) at 24 hours post infection by immunoblotting. (**C**) Assessment of NCOA4 levels in murine macrophages infected with Mtb H37Rv for the indicated time points by immunoblotting. (**D**) Assessment of ferritin heavy chain levels in murine macrophages transfected with NT or *Ncoa4* siRNAs at 24 hours post infection by immunoblotting. (**E**) Confocal images of murine macrophages pretreated with PRMT5 inhibitor (for 1 hour) stained for ferritin heavy chain (green), autophagosomes LC3A/B (red) and nuclear staining (blue) at 24 hours post infection. Scale bars: 2 µm (left), and its quantification (right). (**F**) BALB/c mice were infected with Mtb H37Rv and treated with PRMT5 inhibitor EPZ015666 (4mg/kg) q.o.d. Assessment of ferritin heavy chain levels in lung sections stained for ferritin heavy chain (green) and nuclear staining (blue). The representative confocal image (left) and its quantification (right). All immunofluorescence and immunoblotting data are representative of three independent experiments. ACTB was used as a loading control DAPI, 4',6-diamidino-2-phenylindole; CTCF, corrected total cell fluorescence; Chloroquine (10 µM); MG-132 (10 µM); ****, $p < 0.0001$ (One-way ANOVA in F, Student's t-test in E; GraphPad Prism 10.0). Below each panel, the quantification of the blots normalized to the loading control has been indicated.

**Figure S3.**

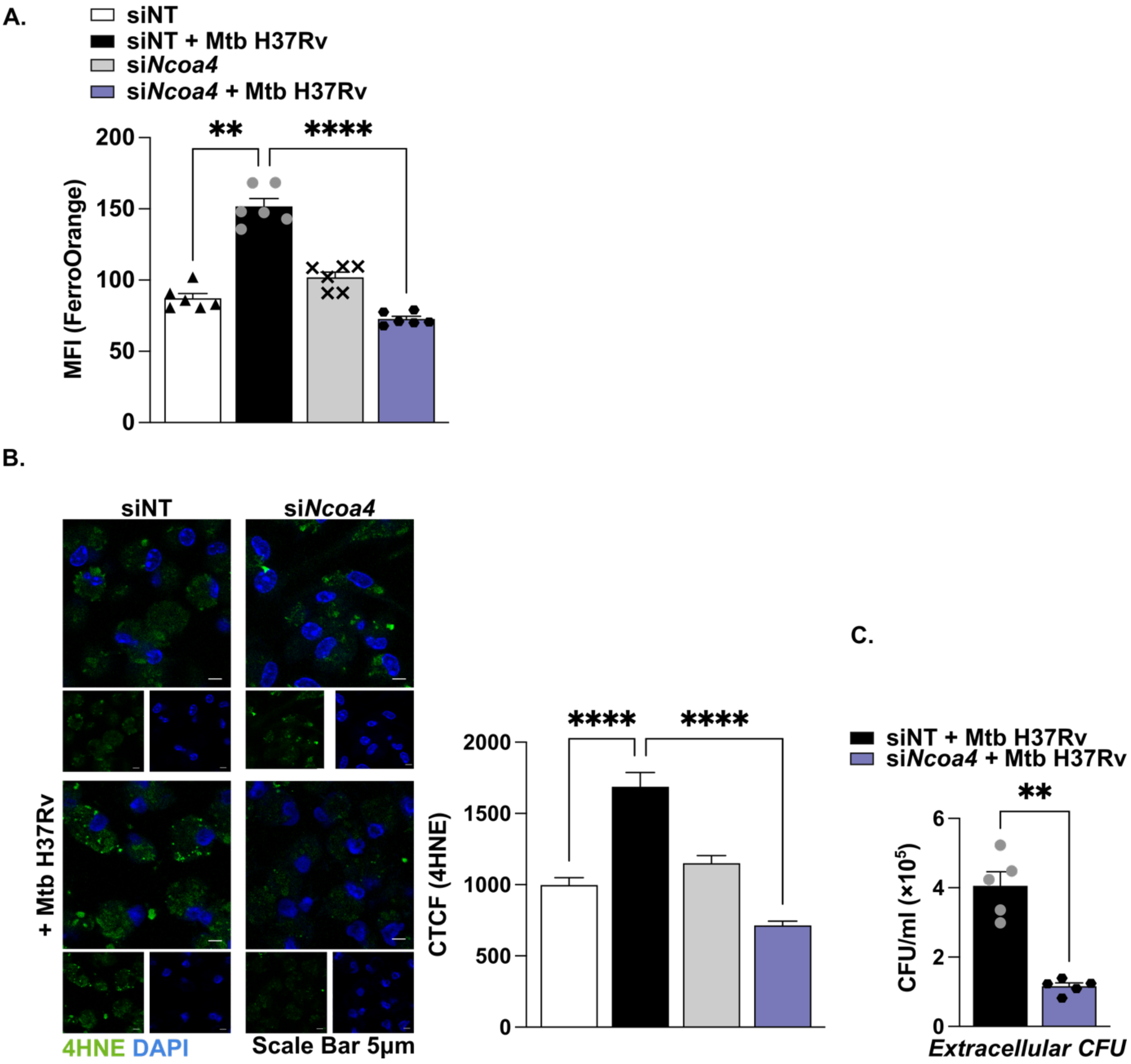

**Figure S3:** (**A-C**) Murine macrophages were transfected with NT or *Ncoa4* siRNAs. (**A**) Estimation of intracellular labile ferrous at 24 hours post infection. (**B**) Confocal images stained for 4HNE (green) and nuclear staining (blue) at 24 hours post infection. Scale bars: 5 µm (left), and its quantification (right). (**C**) Examination of the release of live mycobacteria from necrotic cells by CFU quantification in macrophage culture supernatants at 4 days post infection. All immunoblotting and immunofluorescence data are representative of three independent experiments. NT, non-targeting; MFI, mean fluorescence intensity; DAPI, 4',6-diamidino-2-phenylindole; OE, over expression; CTCF, corrected total cell fluorescence; CFU, colony forming units. **, $p<0.01$; ****, $p < 0.0001$ (One-way ANOVA in A, B; Student's t-test in C; GraphPad Prism 10.0).

**Figure S4.**

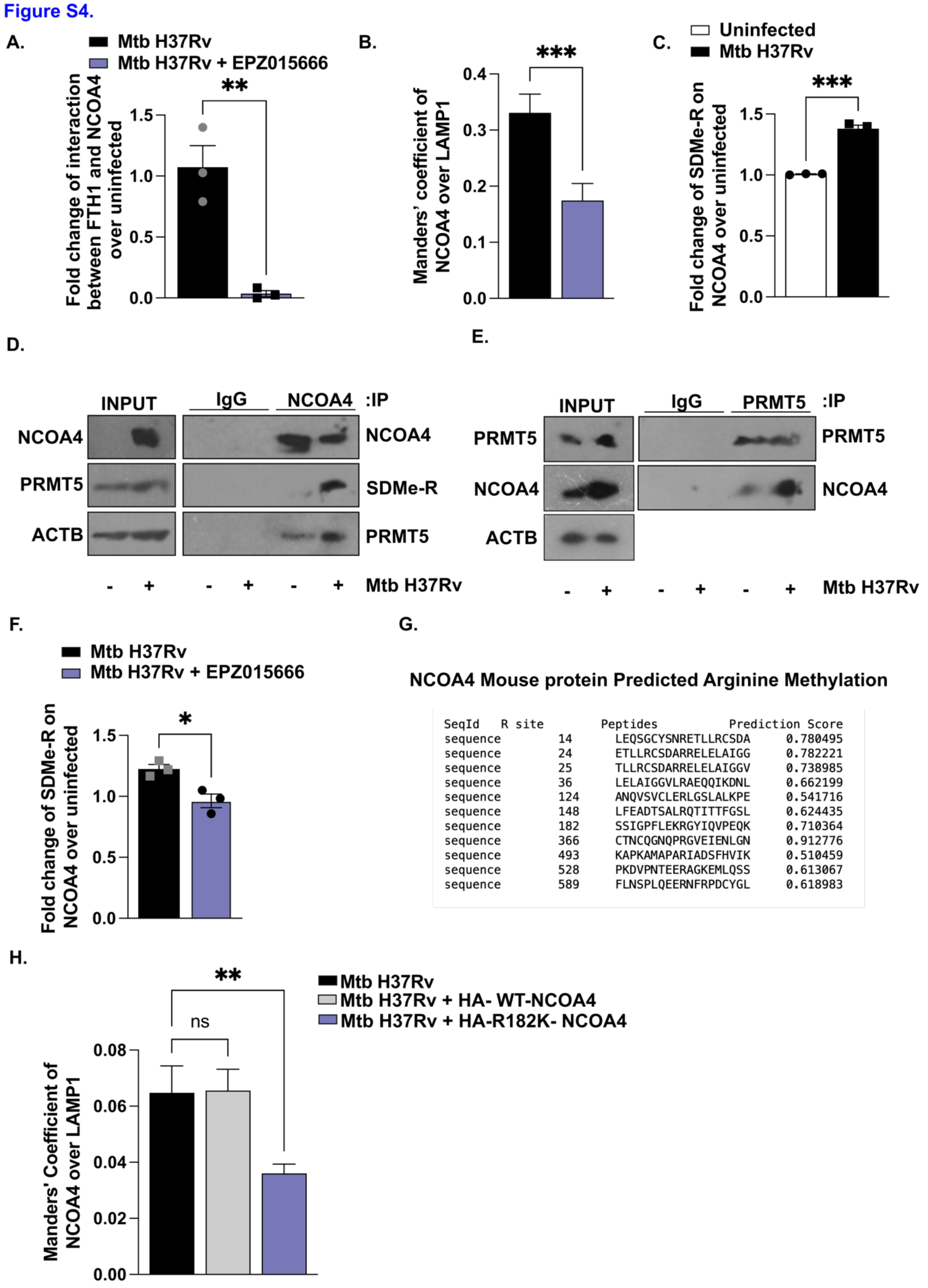


| SeqId | R site | Peptides | Prediction Score |
|---|---|---|---|
| sequence | 14 | LEQSGCYSNRETLLRCSDA | 0.780495 |
| sequence | 24 | ETLLRCSDARRELELAIGG | 0.782221 |
| sequence | 25 | TLLRCSDARRELELAIGGV | 0.738985 |
| sequence | 36 | LELAIGGVLRAEQQIKDNL | 0.662199 |
| sequence | 124 | ANQVSVCLERLGSLALKPE | 0.541716 |
| sequence | 148 | LFEADTSALRQTITTFGSL | 0.624435 |
| sequence | 182 | SSIGPFLEKRGYIQVPEQK | 0.710364 |
| sequence | 366 | CTNCQGNQPRGVEIENLGN | 0.912776 |
| sequence | 493 | KAPKAMAPARIADSFHVIK | 0.510459 |
| sequence | 528 | PKDVPNTEERAGKEMLQSS | 0.613067 |
| sequence | 589 | FLNSPLQEERNFRPDCYGL | 0.618983 |

**Figure S4:** (**A**) Quantification of the interaction between NCOA4 and ferritin heavy chain by co-immunoprecipitation of whole cell lysates from murine macrophages pretreated with PRMT5 inhibitor (for 1 hour) at 24 hours post infection. Representative image is from three independent experiments. (**B**) Quantification of the colocalization between NCOA4 and LAMP1 using immunofluorescence. Representative image in **Figure 3B.** (**C, D**) Assessment of the interaction between NCOA4 and PRMT5 and the extent of arginine methylation on NCOA4 by co-immunoprecipitation of whole cell lysates from murine macrophages at 24 hours post infection (**D**) and the quantification of the methylation. Representative image is from three independent experiments (**C**). (**E**) Assessment of the interaction between PRMT5 and NCOA4 by co-immunoprecipitation of whole cell lysates from murine macrophages at 24 hours post infection (**F**) Quantification of the extent of arginine methylation on NCOA4 by immunoprecipitation of whole cell lysates from murine macrophages pretreated with PRMT5 inhibitor (for 1 hour) at 24 hours post infection. Representative image is from three independent experiments. (**G**) A predictive tool for methylation, PRmePred, was applied to screen and identify the potential residue on mouse NCOA4 that can be methylated PRMT5. (**H**) Quantification of the colocalization between NCOA4 and LAMP1 using immunofluorescence. Representative image in **Figure 4E.** EPZ015666 (20 μM), PRMT5 inhibitor. *, $p<0.05$; **, $p < 0.01$; ***, $p< 0.001$ (Student's t-test in A, B, D, F, H; GraphPad Prism 10.0).

### Figure S5.

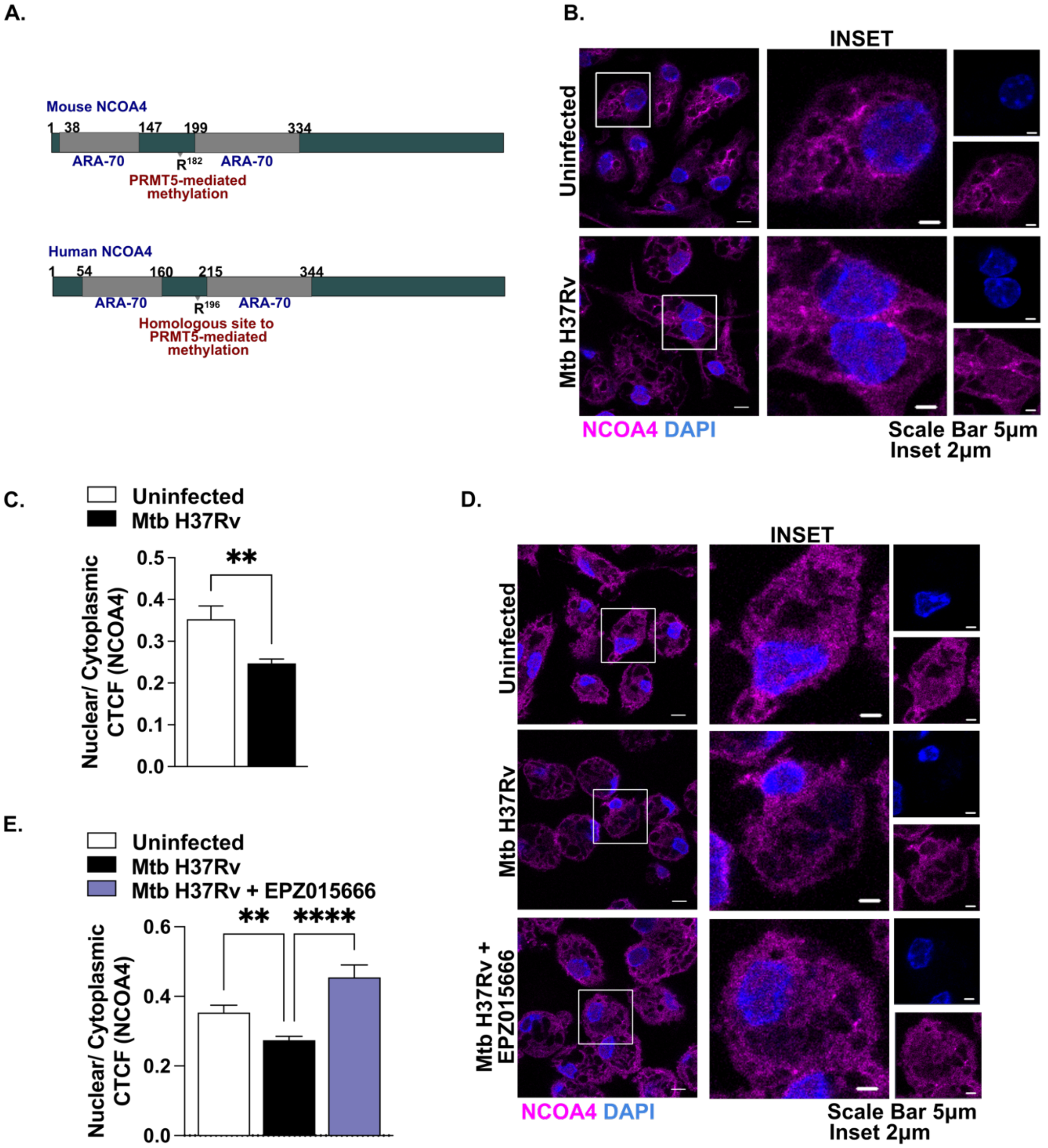


**Figure S5:** (**A**) The methylated arginine residue lies in between two conserved ARA-70 domains in NCOA4 (**B, C**) Confocal images stained for NCOA4 (magenta) and nuclear staining (blue) at 24 hours post infection (**B**) and the quantification of the nuclear cytoplasmic localization (**C**). (**D, E**) Confocal images stained for NCOA4 (magenta) and nuclear staining (blue) in murine macrophages pretreated with PRMT5 inhibitor (for 1 hour) at 24 hours post infection (**D**) and the quantification of the nuclear cytoplasmic localization (**E**). All immunofluorescence data are representative of three independent experiments. EPZ015666 (20 μM), PRMT5 inhibitor. **, p<0.01; ****, $p < 0.0001$ (Student's t-test in C; One-way ANOVA in E; GraphPad Prism 10.0).

**Figure S6.**

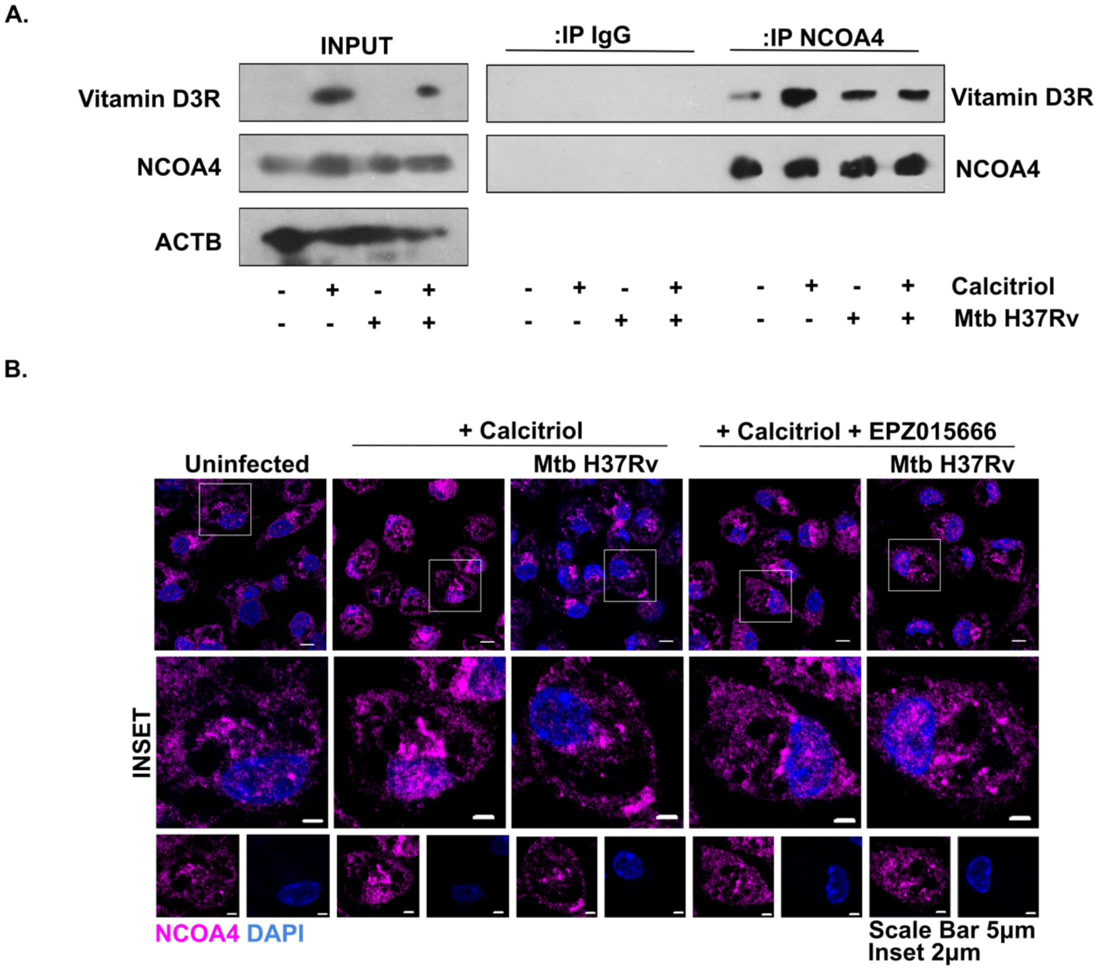


**Figure S6:** (**A**) Assessment of the interaction between NCOA4 and vitamin D3 receptor by co-immunoprecipitation of whole cell lysates from murine macrophages pretreated with Calcitriol (for 1 hour) at 24 hours post infection (**B**) Confocal images stained for NCOA4 (magenta) and nuclear staining (blue) in murine macrophages pretreated with calcitriol and PRMT5 inhibitor (for 1 hour) at 24 hours post infection. All immunofluorescence data are representative of three independent experiments. EPZ015666 (20 μM), PRMT5 inhibitor; Calcitriol (10nM).
